# Genome-wide DNA methylation differences in nucleus accumbens of smokers vs. nonsmokers

**DOI:** 10.1101/781542

**Authors:** Christina A. Markunas, Stephen A. Semick, Bryan C. Quach, Ran Tao, Amy Deep-Soboslay, Laura J. Bierut, Thomas M. Hyde, Joel E. Kleinman, Eric O. Johnson, Andrew E. Jaffe, Dana B. Hancock

**Affiliations:** Center for Omics Discovery and Epidemiology, Behavioral Health Research Division, RTI International, Research Triangle Park, North Carolina; Lieber Institute for Brain Development (LIBD), Baltimore, Maryland; Department of Psychiatry, Washington University School of Medicine, St. Louis, MO; Fellow Program, Behavioral Health Research Division, RTI International, Research Triangle Park, North Carolina

## Abstract

Numerous DNA methylation (DNAm) biomarkers of cigarette smoking have been identified in peripheral blood studies, but their relevance as neurobiological indicators is unknown due to DNAm tissue-specificity. In contrast, blood-based studies may not detect brain-specific smoking-related DNAm differences that may provide greater insight into the neurobiology of smoking behaviors. We report the first epigenome-wide association study (EWAS) of smoking in human postmortem brain, focusing on nucleus accumbens (NAc) as a key brain region in developing addiction. Following Illumina HumanMethylation EPIC array data generation and quality control, 221 decedents (120 European American [23% current smokers], 101 African American [26% current smokers]) were analyzed. DNAm by smoking (current vs. nonsmoking) was tested using robust linear regression models adjusted for age, sex, cell-type proportion, DNAm-derived negative control principal components (PCs), and genotype-derived PCs. Separate ancestry-specific results were combined via meta-analysis, resulting in 7 CpGs that exceeded false discovery rate (FDR)<0.05. Using published smoking EWAS results in blood, we extended our NAc findings to identify DNAm smoking effects that are unique (tissue-specific) versus shared between tissues (tissue-shared). Of the 7 CpGs identified in NAc, 3 CpGs were located near genes previously indicated with blood-based smoking DNAm biomarkers: *ZIC1, ZCCHC24*, and *PRKDC*. The other 4 CpGs are novel for smoking-related DNAm changes: *ABLIM3, APCDD1L, MTMR6*, and *CTCF*. Our results provide the first evidence for smoking-related DNAm changes in human NAc, highlighting CpGs that were previously undetected as peripheral biomarkers and may reflect brain-specific processes.

## INTRODUCTION

Cigarette smoking remains the leading cause of preventable death, resulting in more than 7 million deaths per year worldwide.^1^ Despite the fact that cigarette smoking causes significant morbidity and mortality and nearly 70% of adult smokers report wanting to quit,^2^ 14% of all U.S. adults remain current smokers.^2^

Becoming a regular smoker involves multiple stages, with evidence for heritability at each stage. Heritability estimates range from 37%–55% for smoking initiation,^3, 4^ 46%–59% for smoking persistence,^3^ and up to 75% for nicotine dependence.^4^ Genome-wide association studies (GWAS) have established genetic loci for the varied smoking behaviors (e.g., nicotinic acetylcholine receptor genes),^5^ and, most recently, implicated >400 genetic loci using samples sizes up to 1.2 million individuals.^6^ Despite the strong heritability of smoking behaviors and the successes to date from GWAS, there remains much to be discovered about the neurobiology underlying the trajectory of smoking and the biological mechanisms underlying individual risk loci.^6, 7^

In order to further advance the field and discover novel biological determinants of smoking, alternative study designs, outside of GWAS, are needed. One alternative strategy is to study gene regulation (e.g., epigenetic or gene expression changes associated with disease) in disease-relevant tissues. Brain tissue is highly relevant for studying addiction phenotypes, with different regions implicated in various functions along the addiction reward pathway.^8^ In this study, we focus on the forebrain region, nucleus accumbens (NAc), a key player in reward processes and addiction through its role in cognitive processing of motivation, pleasure, and reward/reinforcement learning.^9, 10^ The NAc influences the dopamine reward system in the first (binge/intoxication) stage of the addiction cycle,^11^ and its disruption has been described as lying at the core of drug addiction.^12^

To date, numerous studies^13-15^ have identified DNA methylation (DNAm) changes related to cigarette smoking in peripheral blood. However, the neurobiological relevance of these changes is unknown due to DNAm tissue-specificity. In addition, these blood-based studies may miss biologically-relevant smoking-related DNAm differences that can only be detected in brain. In this study, we report the first epigenome-wide association study (EWAS) of smoking in human postmortem brain samples, focusing on an addiction-relevant forebrain region, NAc. Using RNA expression (RNAexp) data from the same brain samples, we evaluated smoking-related CpGs for association with nearby RNAexp levels. We also extended our DNAm findings to blood to identify tissue-specific and tissue-shared smoking DNAm effects, and applied a blood-derived smoking DNAm polyepigenetic score^16^ to our brain data to predict current vs. nonsmoking status.

## MATERIALS AND METHODS

### Human post-mortem NAc samples

Postmortem human NAc tissues were obtained at autopsy as part of the Lieber Institute for Brain Development (LIBD) brain collection. Sample exclusion criteria are provided in Supplementary Methods. Information regarding demographics, medical, psychiatric, neurological, and substance use history, as well as current smoking status were collected from a 36-item telephone-administered questionnaire (LIBD Autopsy Questionnaire) with next-of-kin. In addition, cotinine measures from brain and/or blood samples were measured using a standard toxicology screen (3150 B panel; National Medical Services Labs, Inc., Willow Grove, Pennsylvania).

### Tobacco smoke exposure: current smoking vs. nonsmoking

Smoking status was determined based on cotinine levels in blood and/or brain samples and next-of-kin report. Cases (current smokers) were defined by positive cotinine levels (above the reporting limit) in blood and/or brain and a next-of-kin report of current smoking. Controls (nonsmokers) were defined by negative cotinine levels (below the reporting limit) and a next-of-kin report of no current smoking.

### DNAm data, quality control (QC), and pre-processing

Illumina HumanMethylation EPIC data were generated using DNA extracted from NAc samples of 239 eligible decedents, as described previously.^17, 18^ Data quality assessment (Supplementary Methods) and pre-processing were conducted using the R package, *minfi*.^19^ DNAm data processing included stratified quantile normalization, computing principal components (PCs) of the negative control probe intensities to correct for technical artifacts, and estimating neuronal cell type proportions^20^ following the Houseman method^21^ to control for cellular heterogeneity (Supplementary Methods). β-values were calculated to represent the percentage of DNAm at each CpG (ratio of methylated intensities relative to the total intensity).

Quality control resulted in the exclusion of 18 samples and 76,413 probes (Supplementary Methods). The remaining 221 samples and 789,678 probes were analyzed. Sample characteristics are provided in Table 1.

**Table 1.** Description of LIBD NAc samples (N=221)^1^.

| **Variable** | **Case**<br>**(N=53)** | **Control**<br>**(N=168)** | **P-value<sup>2</sup>** |
| --- | --- | --- | --- |
| **Sex, N** |  |  | 0.86 |
| Female | 14 | 42 |  |
| Male | 39 | 126 |  |
| **Race, N** |  |  |  |
| African American | 26 | 75 | 0.57 |
| European American | 27 | 93 |  |
| **Age at death (years), mean ± SD** | 47.25 ± 12.5 | 45.75 ± 13.9 | 0.46 |
| **Proportion of non-neuronal cells<sup>3</sup>,**<br>**mean ± SD** | 0.76 ± 0.07 | 0.76 ± 0.07 | 0.56 |
| **Negative control PCs<sup>4</sup>, mean ± SD** |  |  |  |
| PC1 | -0.08 ± 8.31 | -0.10 ± 11.0 | 0.99 |
| PC2 | 0.06 ± 4.04 | -0.002 ± 3.67 | 0.92 |
| PC3 | -0.06 ± 0.91 | 0.01 ± 2.02 | 0.73 |
| PC4 | -0.15 ± 0.86 | 0.04 ± 1.90 | 0.30 |

<sup>1</sup>Final dataset after quality control and study-specific exclusions were applied to the starting N=239
<sup>2</sup>P-values are based on a Fisher's exact test and t-test for categorical and continuous variables, respectively
<sup>3</sup>Estimated using NAc DNAm data following the Houseman method (refer to Methods for details)
<sup>4</sup>Generated using the negative control probes on the DNAm array (refer to Methods for details)
Frequencies and means ± standard deviations are presented for categorical and continuous variables, respectively
Abbreviations: PC, principal component; SD, standard deviation

### Smoking EWAS meta-analysis in NAc

EWAS was conducted separately by ancestry (European American [EA] and African American [AA] samples) and combined by fixed-effects inverse variance-weighted meta-analysis using *METAL*.^22^ Details regarding model selection are provided in the Supplementary Methods. Robust linear regression analyses were conducted using the R package, *MASS*,^23^ to test the association between current vs. nonsmoking and DNAm (β-value), while adjusting for age at death (continuous), sex (binary), 4 negative control PCs (continuous), genotype PCs (5 for EAs and 10 for AAs), and the estimated proportion of non-neuronal cells. We accounted for multiple testing by controlling the false discovery rate (FDR) at 5%.^24^

An overview of our study design to extend and further characterize our EWAS findings is provided in Figure 1.

**Figure 1.**
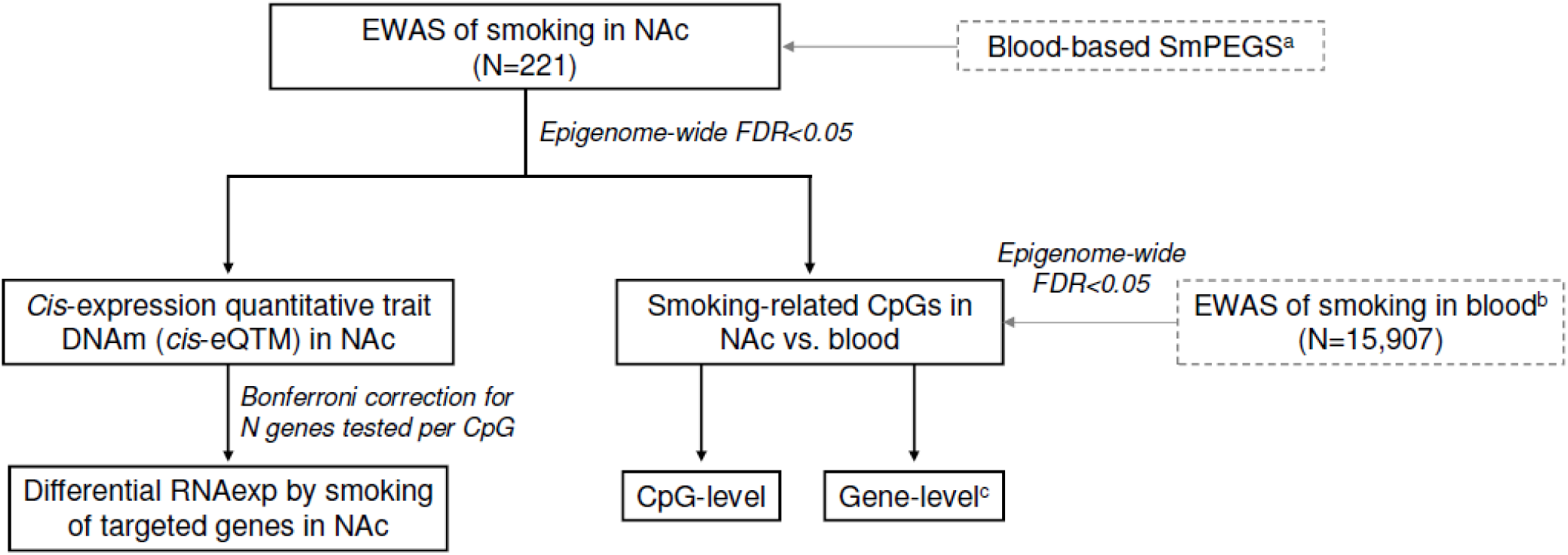
Overview of the study design. Dotted lines indicate previously published results. SmPEGS: Smoking methylation PolyEpigenetic Score. ^a^Sugden K, *et al*. Translational psychiatry 2019. ^b^Joehanes R, *et al*. Circulation Cardiovascular Genetics 2016. ^c^Genes proximal to differentially methylated CpGs.

### DNAm by RNAexp and differential RNAexp by smoking analyses in NAc

We generated corresponding RNA sequencing-based gene expression profiles for each of the NAc samples that had DNAm data and integrated these RNAexp data using a two-step approach (see Supplementary Table S1 for RNAexp sample characteristics). First, we directly correlated the levels of DNAm and nearby expressed genes across samples, regardless of smoking status, to identify those CpGs associated with gene expression (and thus more likely to be functional). Next, we tested genes with a significant RNAexp–DNAm association for differential RNAexp by smoking in NAc, with and without adjustment for nearby DNAm levels. Details regarding RNA-seq data generation, processing, QC, RNAexp–DNAm association analyses, and differential RNAexp by smoking analyses are provided in Supplementary Methods.

### Comparison of smoking-related DNAm changes in NAc and blood

To investigate tissue-specific vs. -shared smoking-related DNAm effects, we compared our EWAS meta-analysis results in NAc (FDR<0.05) to published results from the largest blood-based EWAS meta-analysis of current vs. never smoking to date (N=15,907).^14^ We assessed the overlap between the NAc- and blood-based smoking DNAm results at both the CpG- and gene-level. For the CpG-level comparison, we performed a look-up of our NAc smoking-associated CpGs (FDR<0.05) in the blood-based results, and a reverse look-up of the blood smoking-associated CpGs (FDR<0.05) in our NAc-based results. For the gene-level comparison, we identified genes proximal to smoking-related CpGs that were either specific to NAc or identified in both tissues. Details regarding the CpG- and gene-level comparisons are provided in Supplementary Methods.

### Application of a blood-based Smoking methylation PolyEpigenetic Score (SmPEGS) in brain

We applied the blood-based SmPEGS, as developed by Sugden^16^ *et al.* to our brain DNAm data (Supplementary Methods). Logistic regression models were run separately by ancestry and in the pooled sample to test the association between the blood-derived SmPEGS and current vs. nonsmoking in NAc, adjusting for the same covariates used in our brain EWAS analysis. To assess the ability of the SmPEGS to classify smoking status in brain, we performed a receiver operating characteristic (ROC) analysis using the R package, *pROC* (Supplementary Methods).^25^

## RESULTS

### Epigenome-wide association results by smoking in NAc

We identified 7 CpGs that exceeded FDR<0.05 in our cross-ancestry EWAS meta-analysis (Figure 2 and Table 2; Supplementary Figures S1 and S2). The difference in the mean percentage of DNAm in smokers and nonsmokers ranged from 0.92% to 4.13% for these 7 CpGs. Three of the 7 smoking-associated CpGs exhibited decreased DNAm by smoking, with consistent directions of effect between EA and AA ancestries (Table 2). Overall, smoking-related effect sizes were small, and there was no evidence of systematically increased or decreased DNAm by smoking (Figure S1). More complete EWAS results are provided in Supplementary Table S2 (Meta-analysis P<0.05) and full results are provided in Supplementary Table S3.

**Table 2.**
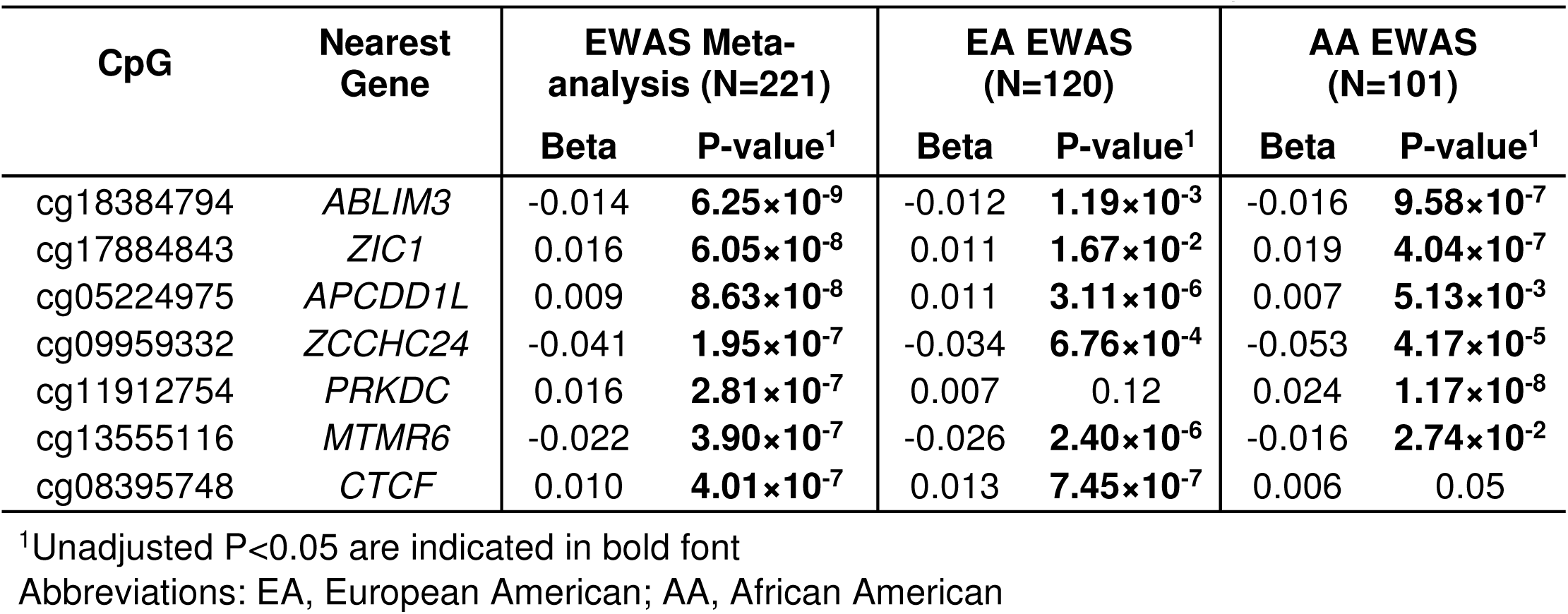
Smoking-related CpGs in NAc at FDR<0.05 in EWAS meta-analysis.

**Figure 2.**
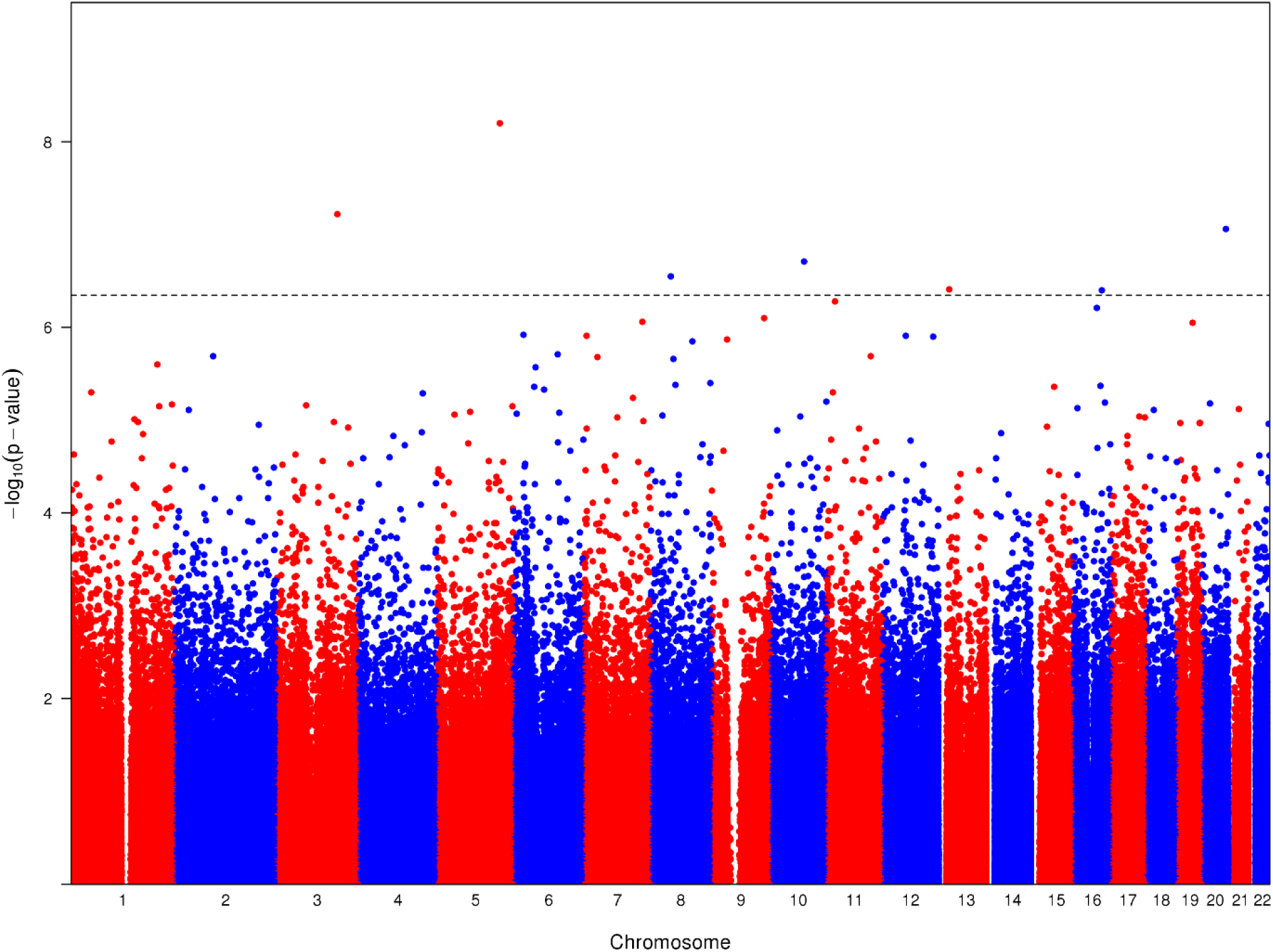
EWAS meta-analysis in NAc of smokers (N=53) vs. nonsmokers (N=168). CpGs are shown according to their position on chromosomes 1–22 (alternating red/blue) and plotted against their -log_10_ p-values. The dotted horizontal line indicates genome-wide significance based on FDR<0.05. The genomic inflation factor (λ) = 1.05 for the meta-analysis and 1.01 and 0.98 for European American- and African American-specific analyses, respectively.

### Linking DNAm associations with smoking to RNAexp in NAc

Using a two-step approach, we integrated DNAm data with corresponding RNAexp data in the same NAc samples to identify: (1) smoking-related CpGs associated with nearby RNAexp, and (2) genes with differential RNAexp by smoking. We first focused on the 7 smoking-related CpGs in NAc, and tested for association between DNAm levels and RNAexp of each gene within a 1 Mb interval (Supplementary Table S4). We identified one long noncoding RNA (lncRNA) with RNAexp levels that were significantly associated with DNAm levels at cg08395748: *RP11-297D21.4* (ENSG00000270049: antisense to *AGRP* [Agouti related neuropeptide]; P=1.54×10^−4^, located 0.07 Mb upstream). We next evaluated *RP11-297D21.4* for differential RNAexp by smoking and found no evidence of an association, regardless of controlling for nearby DNAm levels at cg08395748 (Without adjustment for DNAm: log_2_ fold change=-0.06, P=0.16; With adjustment for DNAm: log_2_ fold change=-0.03, P=0.60). Overall, these results provide limited functional support for the smoking-related DNAm changes in NAc. Out of the 7 smoking-related CpGs, only cg08395748 was associated with nearby RNAexp (*RP11-297D21.4*), but further evaluation of *RP11-297D21.4* indicated that it was not differentially expressed by smoking.

### Comparisons to infer tissue-specific and -shared smoking DNAm effects

#### CpG-level: Comparison of differentially methylated CpGs

To assess whether our NAc findings are present in peripheral tissues outside the brain, we used the largest reported differential DNAm study by smoking in blood^14^ using the Illumina 450k array and found that none of the 5 present smoking-related CpGs in NAc were associated in blood (Table 3). The 2 other smoking-related CpGs in NAc were captured on the EPIC array, but not the 450K array, and thus were not tested in blood. The smoking effect sizes for all 5 of the available smoking-related CpGs in NAc were smaller in blood than in brain, and the directions of effect were consistent for 3 of the 5 CpGs. The mean percentages of DNAm in blood and brain were similar.

**Table 3.**
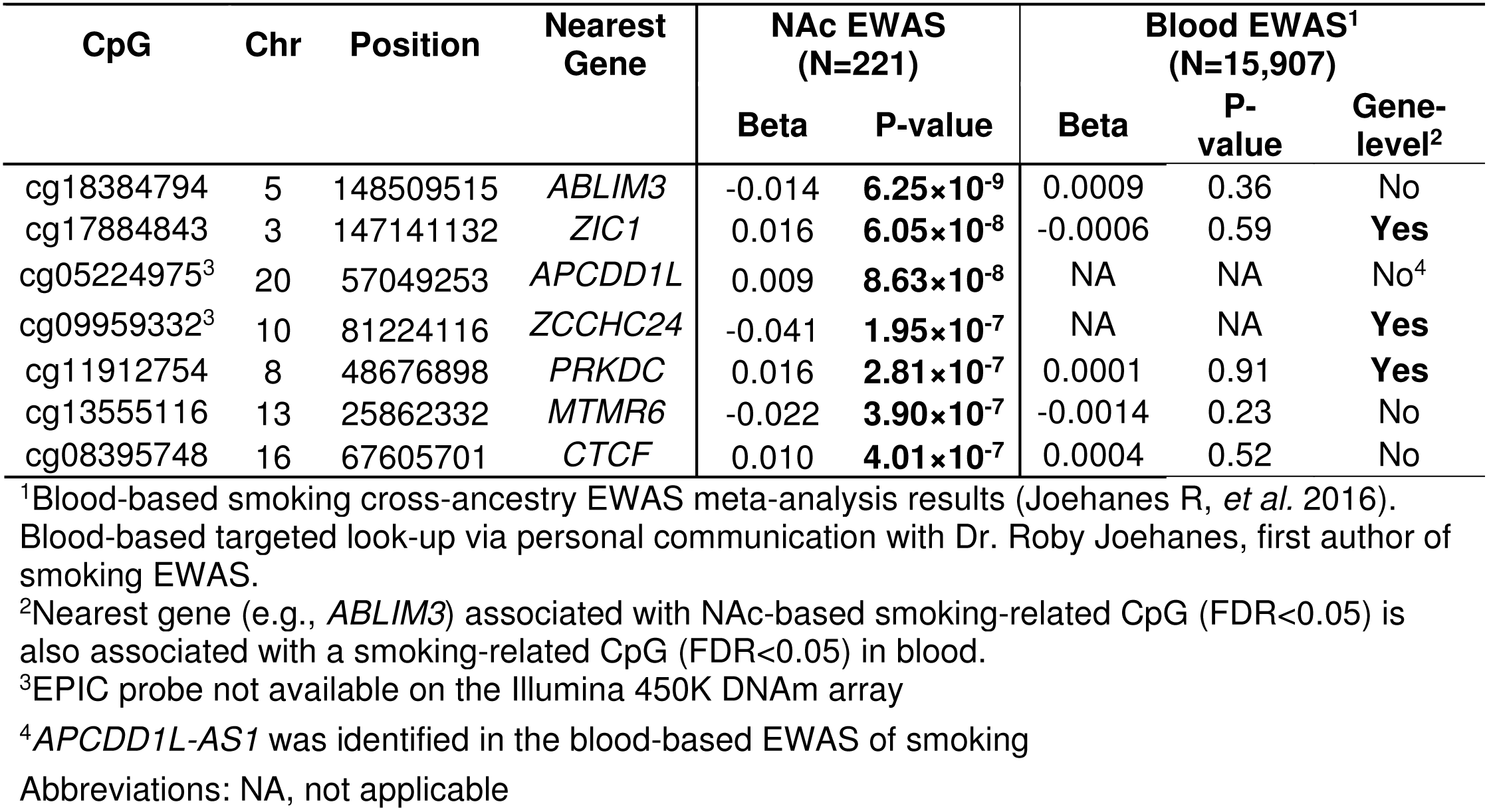
Overlap between smoking-related CpGs in NAc and blood samples (FDR<0.05).

We also performed the reverse look-up of blood-based findings in NAc. None of the 16,706 present smoking-related CpGs in blood were associated in NAc (Supplementary Table S5) and the effect sizes were not correlated (r=0.08). Overall, we found no evidence of tissue-shared smoking-related DNAm changes at the CpG-level.

#### Gene-level: Comparison of genes proximal to differentially methylated CpGs

We annotated the 7 CpGs with differential DNAm by smoking in NAc to the nearest gene, and among the 7 unique genes identified, 3 genes (*ZIC1* [Zic family member 1], *ZCCHC24* [Zinc finger CCHC-type containing 24], *PRKDC* [Protein kinase, DNA-activated, catalytic subunit]) were also annotated as the nearest gene to a CpG that showed evidence of differential DNAm by smoking in blood (Table 3 and Supplementary Table S6). Among the 3 overlapping genes, *ZIC1* was the only gene for which the smoking-associated CpG in NAc fell within the gene body. While the specific CpG sites with evidence of DNAm changes by smoking were not shared between tissues, the directions of effect for associated CpGs at FDR<0.05 were consistent between tissues for *ZIC1* (higher DNAm levels among smokers) and *ZCCHC24* (lower DNAm levels among smokers), but not for *PRKDC*. Overall, our results support both tissue-specific (*ABLIM3* [Actin binding LIM protein family member 3], *APCDD1L* [APC down-regulated 1 like], *MTMR6* [Myotubularin related protein 6], and *CTCF* [CCCTC-binding factor]) and -shared (*ZIC1, ZCCHC24, PRKDC*) genes proximal to differentially methylated CpG(s) by smoking.

### Blood-based SmPEGS tests for discrimination of smoking status in brain

We applied the blood-based SmPEGS^16^ to our brain DNAm data to predict smoking status (Supplementary Table S7). The SmPEGS was associated with smoking in the combined (OR=1.78 [95% confidence interval: 1.22, 2.59], P=0.003) and EA (OR=2.00 [1.12, 3.59], P=0.02) samples. The association among AAs was non-significant but in the same direction (OR=1.45 [0.79, 2.67], P=0.24). The blood-based SmPEGS provided limited discrimination between smokers and nonsmokers in Nac with an area under the ROC curve (AUC)=0.62 (95% confidence interval: 0.54, 0.71) in the combined sample and an AUC=0.67 (0.55, 0.78) in EA samples.

## DISCUSSION

Our results strongly support tissue-specific smoking DNAm effects at the CpG-level in blood versus brain, with some support for tissue-shared DNAm effects at the gene-level. Our study is the first to report epigenome-wide DNAm changes by smoking in human postmortem brain, identifying 7 CpGs exceeding FDR<0.05 in NAc. We identified the smoking-associated cg08395748 as a *cis*-expression quantitative trait DNAm for a lncRNA (*RP11-297D21*) in NAc; however, we found no evidence of differential RNAexp by smoking for *RP11-297D21*. To investigate both tissue-specific and -shared smoking DNAm effects, we extended our NAc DNAm findings to blood. None of the available NAc- or blood-smoking-associated CpGs were associated with smoking in the other tissue. However, 3 of the 7 NAc CpGs were nearby genes that were also identified as proximal to smoking-associated CpGs in the prior blood-based EWAS.

The 3 genes proximal to smoking-related DNAm changes in both blood and NAc were *ZIC1, ZCCHC24*, and *PRKDC*. While different CpGs were implicated in NAc vs. blood, the directions of effect for DNAm levels by smoking were consistent between tissues for *ZIC1* and *ZCCHC24.* All genes have been previously related to smoking in other contexts. Zic1 is a zinc finger transcription factor that plays a role in early development and formation of the cerebellum.^26^ All 5 smoking-associated *ZIC1* CpGs in blood and NAc (FDR<0.05) showed increased DNAm in current smokers. However, the opposite direction of effect has been reported for sperm samples (N=156), with decreased DNAm at *ZIC1* in current smokers (P=9.6×10^−8^).^27^ *ZIC1* has also been implicated in a smoking initiation GWAS meta-analysis of 1.2 million individuals (PASCAL gene-based test: P=4.14×10^−7^).^6^ In addition, dysregulation of *ZIC1* in brain has been observed for another drug exposure, as *ZIC1* is downregulated in response to repeated cocaine administration in mouse NAc (P=3.7×10^−7^).^28^ Taken together, there are multiple lines of evidence linking *ZIC1* to smoking, and possibly drug use more broadly.

Little is known about the function of *ZCCHC24*, but it has been related to smoking previously. Increased DNAm at *ZCCHC24* in newborn blood samples has been associated with sustained maternal smoking during pregnancy (P=5.29×10^−4^).^15^ *PRKDC* encodes the DNA-dependent protein kinase catalytic subunit and plays a key role in the DNA repair pathway, non-homologous end-joining (NHEJ).^29^ Genetic variants in NHEJ genes,^30^ including *PRKDC*,^31^ have been previously associated with lung cancer, of which cigarette smoking is a major risk factor.

In addition to 3 tissue-shared smoking DNAm effects at the gene-level (i.e., genes that are proximal to differentially methylated CpG[s] by smoking), we also report 4 genes with evidence of proximal differential DNAm by smoking only in NAc: (1) *ABLIM3*, (2) *APCDD1L*, (3) *MTMR6*, and (4) *CTCF*. To our knowledge, none of these genes have been previously shown to have DNAm changes by smoking; however, prior links to smoking and/or neurobiology have been demonstrated. *ABLIM3* is involved in axonal guidance signaling and is downregulated in dopaminergic neurons in the ventral tegmental area in relation to perinatal nicotine exposure in rats.^32^ Little is known about the function of *APCDD1L*. Among other genes, *APCDD1L* was selected as part of two blood-based gene expression signatures developed to classify smokers versus nonsmokers.^33^ While *APCDD1L* itself was not annotated to any smoking-associated CpGs in blood, the antisense non-coding RNA of *APCDD1L* (*APCDD1L-AS1*) was identified (cg14546153: β=0.003, P=0.0004).^14^ *MTMR6* is a member of the myotubularin-related protein family of phosphatidylinositol 3-phosphatases and plays a key role in 3-phosphoinositide lipid metabolism.^34^ While *MTMR6* dysregulation in brain hasn’t been previously linked to smoking or addiction phenotypes, differential *MTMR6* expression in brain has been implicated in schizophrenia,^35^ which is highly comorbid and genetically correlated with smoking.^36^ *CTCF* is a well-known transcriptional regulator and thought to play a role in brain development and regulation of neural genes.^37^ Furthermore, it is thought to be involved in changes to astrocyte morphology and function in response to dopamine, a neurotransmitter important in addiction and reward pathways.^38^

Only tissue-specific effects were observed at the CpG-level. This observed tissue specificity by smoke exposure was not restricted to only NAc-identified smoking-related CpGs as we also didn’t identify associations of blood-based smoking-associated CpGs in NAc. For example, cg05575921, located in *AHRR* (Aryl-Hydrocarbon Receptor Repressor), is one of the most robustly replicated blood-based DNAm biomarkers for cigarette smoking in adults, but was not associated with smoking in NAc (Meta-analysis P=0.57; 0.19% difference in DNAm between smokers and nonsmokers). For comparison, cg05575921 showed an 18% difference in DNAm between smokers and never smokers in blood samples (P=4.5×10^−26^).^14^ While unlikely to fully account for the lack of association in brain, the likely presence of former smokers among our nonsmoker controls may reduce our power to detect DNAm changes at cg05575921, as cg05575921 is associated with former vs. never smoking, although the effect is attenuated by 77.5% (as compared to current vs. never smokers).^14^

To further assess how well blood-based findings extend to brain, we applied a blood-derived smoking DNAm polyepigenetic score to our brain data to predict smoking status. We found that the SmPEGS provided limited discrimination between smokers and nonsmokers in NAc (AUC=0.62 [0.54, 0.71]), indicating that many of the effects detected in blood may not directly translate to strong effects in brain. For comparison, AUCs of 0.81 (E-Risk Longitudinal Twin Study [E-Risk]) and 0.93 (Dunedin Longitudinal Study [Dunedin]) for never vs. current smoking and 0.78 (E-Risk) and 0.77 (Dunedin) for never vs. ever (current and former smokers) smoking were reported in blood samples from the original SmPEGS publication.^16^ These AUCs were derived from samples not included as part of the blood-based smoking EWAS meta-analysis,^14^ from which the SmPEGS was derived.^16^

While the data from this study represent a powerful resource to begin to investigate the neurobiological underpinnings of nicotine use disorder and other smoking phenotypes, there are several limitations. Due to the highly unique nature of the data (i.e., multi-omic data in human postmortem brain with available smoking data), our sample size and thus statistical power was limited which could impact our ability to detect smoking-related DNAm changes in brain and link those changes with nearby RNAexp. Further, there are currently no external NAc DNAm datasets for replication of our novel findings. We therefore relied on internal consistency between ancestral groups by focusing on meta-analysis results, rather than ancestry-specific findings. In addition, comparisons to blood may have been affected by differences in the statistical power between datasets and/or the type of Illumina DNAm array used. While the power to detect blood-based findings in brain is more limited, there should be sufficient power to detect the NAc-based findings in the blood (N=15,907). Although different Illumina DNAm arrays were used (Illumina 450K [Blood] vs. EPIC [NAc] array), the more recently developed Illumina EPIC array targets over 90% of the 450K CpG probes and both arrays use the same probe chemistries and generally exhibit high levels of correlation.^39^ In addition, the reference data used to derive estimates of cell type proportions were based on Illumina 450K data generated in dorsolateral prefrontal cortex brain samples.^20^ While we used cell type-discriminating CpGs that overlapped with the EPIC array for cell type proportion estimates in NAc, there may be residual cell type differences that were unaccounted for in our analysis. Finally, we are unable to rule out residual confounding due to other exposures or traits (e.g., current/former alcohol use [no decedent was intoxicated at time of death]) or reduced power due to smoking-related DNAm changes which persist in former smokers.

Our study is the first to demonstrate smoking-related DNAm changes in human brain and lays the foundation for future studies to expand and extend our findings in brain. Our results highlight CpGs that were previously undetected as peripheral DNAm biomarkers of smoking and may reflect brain-specific processes, as well as provide support for tissue-shared genes proximal to smoking-related DNAm changes, suggesting that the susceptibility of certain epigenomic regions to smoking may be conserved between tissues. Importantly, the DNAm changes identified in this study may underlie neurobiological processes related to smoke exposure and/or reflect a genetic predisposition for smoking. Future work will involve incorporating genetic data to begin to differentiate smoking-related DNAm changes that are genetically-driven (i.e., DNAm as a mediator of genetic risk of smoking) versus exposure-driven (i.e., DNAm changes as a consequence of exposure). We expect that increasing sample size and expanding the number of brain tissues examined in future studies will identify additional DNAm and RNAexp changes and enable the investigation of ancestry-specific effects.

## ACKNOWLEDGEMENTS

This work was supported by the National Institute on Drug Abuse R01 DA042090 (PI: Hancock).

