## Supplementary Materials for "Genome-wide DNA methylation differences in nucleus accumbens of smokers vs. nonsmokers"

### Supplementary Methods:

#### Human post-mortem NAc samples

All decedents included in this study had a postmortem interval less than 72 hours. Decedents with brain trauma, metastatic brain cancer, neuritic pathology, neurodegenerative diseases, HIV/AIDS, hepatitis, or other communicable diseases were excluded. In addition, all decedents were reviewed by a board-certified psychiatrist. Decedents determined to have a DSM-5 lifetime psychiatric or substance use disorder diagnosis, with the exception of nicotine use disorder, were excluded. Analyses were restricted to decedents with a minimum age at death of 13 years old. Ethanol toxicology testing was completed on every donor and none of the decedents selected for our study had blood alcohol levels over the legal limit.

#### DNAm data, quality control (QC), and pre-processing

Overall data quality was assessed using a series of diagnostic plots to identify outlier samples. Samples were checked for sex and race mismatches, low call rate (≥ 1% of probes failed [Illumina detection P>0.01]), and sample swaps. Sample swaps were determined based on pairwise genotype correlations (Pearson's r) calculated from 1000 Genomes Phase 3 genotype data imputed from Illumina arrays and DNAm array data (using SNP probes) on the same set of samples. Problematic samples were defined as having r>0.8 for non-matching sample IDs and r<0.7 for matching sample IDs.

We determined which estimated cell type proportions (neuronal cells, non-neuronal cells, embryonic stem cells, neural progenitor cells, and dopaminergic neurons) to include in the final model based on the percentage of explained variance of the first principal component (PC1) of DNAm, which largely represents variation due to cellular heterogeneity. When all cell types were included in the same model (DNAm PC1 ~ neuronal cells + non-neuronal cells + embryonic stem cells + neural progenitor cells + dopaminergic neurons), only the proportion of neuronal cells, non-neuronal cells, and embryonic stem cells were significantly associated ($P<0.05$) with DNAm PC1. As the estimated proportion of neuronal cells was highly correlated with the proportion of non-neuronal cells ($r=0.97$), we dropped neuronal cells from the model as non-neuronal cells alone explained more of the variance of DNAm PC1. The proportion of non-neuronal cells explained 95.8% of variation of DNAm PC1, compared with neuronal cells explaining 93.6% of variation, and was selected for inclusion in the final model, as only slight improvements in the percentage of variance explained was achieved by inclusion of embryonic stem cells as well (0.16% difference in variance explained).

Sample and probe-level exclusions were applied to the data. In total, 18 samples were excluded due to cross- and within-data genotype QC checks (N=11), indeterminate smoking status (N=2), missing genotype data (N=2), and outlier based on PCA plot of DNAm beta values (N=3). Probes that failed in over 5% of samples (N=2,118), contained a common (MAF>1%) SNP (N=14,407) at the single base extension site, did not uniquely map to the genome[1] (N=42,126), or were located on a sex chromosome (N=17,726) were excluded.

**Smoking epigenome-wide association study (EWAS) meta-analysis in NAc**

A series of sensitivity analyses were conducted to determine a final model with respect to the number of negative control PCs and genotype PCs. The most parsimonious model that achieved a genomic inflation factor (λ) near 1 was selected (final model described in main manuscript).

**DNAm by RNAexp and differential RNAexp by smoking analyses in NAc**

*RNA sequencing (RNA-seq) and processing*

The same set of Lieber Institute for Brain Development (LIBD) postmortem NAc samples used for DNAm analysis were processed and prepared for RNA-seq, as described previously for dorsolateral prefrontal cortex brain samples.[2] A total of 239 RNA samples were sequenced using paired-end 100 bp reads on an Illumina HiSeq3000 at LIBD. Following RNA-seq, samples were first checked for read adapter content to identify adapter contamination. No samples failed the adapter content check, thus reads were not trimmed prior to mapping. Reads were mapped to the GRCh38 reference transcriptome (GENCODE v25) using *HISAT2,*[3] and transcript-level quantifications were estimated using *Salmon*.[4] Gene-level quantifications for downstream differential RNAexp analysis were derived from the transcript-level estimates using the R package, *tximport*.[5]

*RNA-seq data QC*

To identify potential sample swaps, genotype correlations derived from 1000 Genomes Phase 3 imputed genotype and RNA-seq data were calculated. Genotypes were called from RNA-seq data using the *mpileup* utility from *SAMTools*.[6] Problematic samples were identified by calculating pairwise genotype correlations (Pearson's *r*) within tissue, as well as between RNA-seq and genotype datasets. Samples were either excluded or mismatches resolved if r>0.8 for non-matching sample IDs or r<0.7 for matching sample IDs.

In addition to identifying sample swaps, sex discrepancies were assessed. Males and females formed two distinct chromosome Y gene expression clusters, resulting in a bimodal distribution of per-sample mean counts per million (CPM) values across chromosome Y genes. Consequently, discrepant sex assignments based on k-means (k=2) clustering of the mean CPM values of chromosome Y genes were excluded. The following sample-level exclusions were applied (N=40 excluded; exclusion criteria are not mutually exclusive): RIN<6 (N=16), overall mapping rate <0.50 or gene assignment rate <0.30 or mitochondrial mapping rate >0.11 (N=6); missing phenotype data (N=2); missing genotype data (N=4); problematic based on genotype correlations (N=20).

*RNAexp–DNAm association analysis in NAc*

We tested for association between DNAm levels and RNAexp of each gene within a 1 Mb interval (using the same GRCh38 reference transcriptome [GENCODE v25] used for RNA-seq read mapping) of the smoking-associated CpGs (FDR<0.05). RNAexp–DNAm association analyses were conducted using 196 samples with both DNAm and RNA-seq QC'd data available, following the model below (**Equation 1**).

$$RNAexp_i = \alpha_i + \beta_1 DNAm_i + \beta_2 age_i + \beta_3 sex_i + \beta_4 RIN_i + \beta_5 chrM_i + \beta_6 GAR_i + \beta_7 GenotypePC_i^1 + \cdots + \beta_{11} GenotypePC_i^5 + \beta_{12} RNAexpSV_i^1 + \cdots + \beta_{30} RNAexpSV_i^{19} + \beta_{31} DNAmPC_i^1 + \cdots + \beta_{34} DNAmPC_i^4 + \beta_{35} CellProp_i + \epsilon_i \quad \textbf{(1)}$$

Here $RNAexp_i$ represents the $\log_2$ transformed, median-ratio normalized (across sample normalization performed by DESeq2[7]) gene counts, $DNAm_i$ represents the

DNAm β-values, $age_i$ is age at death, $sex_i$ is a binary variable for sex, $RIN_i$ is RNA integrity number, $chrM_i$ is mitochondrial read mapping rate, $GAR_i$ is gene assignment rate, $GenotypePC_i^{1-5}$ corresponds to the top five principal components calculated using LD-pruned genotyping array data, $RNAexpSV_i^{1-19}$ corresponds to the 19 significant surrogate variables estimated using *SVA*[8], $DNAmPC_i^{1-4}$ corresponds to the 4 negative control DNAm PCs, and $CellProp_i$ refers to the estimated proportion of non-neuronal cells. Prior to model fitting, genes that did not have more than 10 counts in greater than 10% of the samples were removed. A Bonferroni correction was applied to control for the number of genes tested per CpG.

*Differential RNAexp by smoking analysis*

Genes with significant RNAexp–DNAm associations were next tested for differential gene-level RNAexp by smoking in NAc, with (**Equation 2**) and without (**Equation 3**) adjustment for DNAm at the *cis*-expression quantitative trait DNAm (*cis*-eQTM). Differential gene expression analyses were conducted on 196 samples with both DNAm and RNA-seq QC'd data available using *DESeq2*[7] with the following models for each gene$i$:

$$y_i = \alpha_i + \beta_1 smoking_i + \beta_2 DNAm_i + \beta_3 age_i + \beta_4 sex_i + \beta_5 RIN_i + \beta_6 chrM_i + \beta_7 GAR_i + \beta_8 GenotypePC_i^1 + \cdots + \beta_{12} GenotypePC_i^5 + \beta_{13} RNAexpSV_i^1 + \cdots + \beta_{31} RNAexpSV_i^{19} + \beta_{32} DNAmPC_i^1 + \cdots + \beta_{35} DNAmPC_i^4 + \beta_{36} CellProp_i + \epsilon_i \quad \textbf{(2)}$$

$$y_i = \alpha_i + \beta_1 smoking_i + \beta_2 age_i + \beta_3 sex_i + \beta_4 RIN_i + \beta_5 chrM_i + \beta_6 GAR_i + \beta_7 GenotypePC_i^1 + \cdots + \beta_{11} GenotypePC_i^5 + \beta_{12} RNAexpSV_i^1 + \cdots + \beta_{30} RNAexpSV_i^{19} + \epsilon_i \quad \textbf{(3)}$$

Here $smoking_i$ indicates case/control smoking status (as defined for DNAm analyses), $DNAm_i$ represents the DNAm β-values at the *cis*-eQTM, $age_i$ is age at death, $sex_i$ is a binary variable for sex, $RIN_i$ is RNA integrity number, $chrM_i$ is mitochondrial read mapping rate, $GAR_i$ is gene assignment rate, $GenotypePC_i^{1-5}$ corresponds to the top five principal components calculated using LD-pruned genotyping array data, $RNAexpSV_i^{1-19}$ corresponds to the 19 significant surrogate variables estimated using *SVA*[8], $DNAmPC_i^{1-4}$ corresponds to the 4 negative control DNAm PCs, and $CellProp_i$ refers to the estimated proportion of non-neuronal cells.

**Comparison of smoking-related DNAm changes in NAc and blood**

To investigate tissue-specific vs. -shared smoking-related DNAm effects, we compared our EWAS meta-analysis results in NAc (FDR<0.05) to published results from the largest blood-based EWAS meta-analysis of smoking to date (N=15,907).[9] Specifically, we used the 18,760 epigenome-wide significant results (FDR<0.05; published Supplementary Table 2[9]) from the current vs. never smoking cross-ancestry EWAS meta-analysis based on Illumina 450K DNAm data. We assessed the overlap between our NAc- and blood-based smoking DNAm results at both the CpG- and gene-level. For the CpG-level comparison, we performed a look-up of our NAc smoking-associated CpGs (FDR<0.05) in the blood-based results, and a reverse look-up of the blood smoking-associated CpGs (FDR<0.05) in our NAc-based results. A Bonferroni correction was applied to control for the number of CpGs tested. For the gene-level comparison, we identified genes proximal to smoking-related CpGs that were either specific to NAc or identified in both tissues. We used *BEDTools*[10] to assign each CpG to the closest GENCODE v29 gene (GRCh37/hg19) within a 1 Mb interval. When a CpG overlapped >1 gene, all genes were annotated to the CpG. For each set of smoking-associated CpGs (blood-based and brain-based FDR<0.05 results), a list of unique genes was created for comparison.

**Application of a blood-based Smoking methylation PolyEpigenetic Score (SmPEGS) in brain tissue**

To calculate the SmPEGS, we followed the approach developed by Sugden et al.[11] The SmPEGS[11] was derived using the 2,623 CpGs identified as significantly associated with current vs. never smoking ($P<1\times10^{-7}$) in the blood-based cross-ancestry EWAS meta-analysis[9] that were also available in our brain data. Using the overlapping probes (2,345 present in NAc QC'd data) in our brain data, we multiplied the DNAm β-value by the regression coefficient estimated in the blood-based smoking EWAS meta-analysis.[9] The SmPEGS was computed as the mean of the weighted β-values (DNAm β-value in NAc × blood-based regression coefficient) across the set of 2,345 CpGs and then standardized ($\mu=0$, $\sigma=1$) across individuals. These SmPEGSs were used in logistic regression models to test the association between the blood-derived SmPEGS and current vs. nonsmoking in NAc, adjusting for the same covariates used in our brain EWAS analysis.

To assess the ability of the SmPEGS to classify smoking status (current smoking vs. nonsmoking) in brain, we performed a receiver operating characteristic (ROC) analysis using the R package, *pROC*. [12] To use as input into the ROC analysis, we regressed SmPEGS values on the same set of covariates used in the brain EWAS analysis, obtained residuals, and standardized ($\mu=0$, $\sigma=1$) within the sample. The 95%

confidence interval of the area under the ROC curve (AUC) was calculated using 5,000 bootstrap iterations.

### Supplementary Results:

**Table S1. Description of LIBD NAc RNA-seq samples (N=199)[1].**

| Variable | Case (N=46) | Control (N=153) | P-value[2] |
|---|---|---|---|
| **Sex, N** | | | 0.44 |
| Female | 14 | 37 | |
| Male | 32 | 116 | |
| **Race, N** | | | 0.74 |
| African American | 23 | 71 | |
| European American | 23 | 82 | |
| **Age at death (years), mean ± SD** | 48.59 ± 11.69 | 45.58 ± 14.21 | 0.15 |
| **RNA integrity number, mean ± SD** | 8.00 ± 1.05 | 7.52 ± 0.46 | **$4.0\times10^{-3}$** |
| **Mitochondrial read mapping rate, mean ± SD** | 0.04 ± 0.02 | 0.04 ± 0.02 | 0.60 |
| **Gene assignment rate, mean ± SD** | 0.44 ± 0.04 | 0.45 ± 0.05 | **$2.0\times10^{-2}$** |

[1]Final dataset after quality control and study-specific exclusions were applied

Frequencies and means are presented for categorical and continuous variables, respectively

[2]P-values are based on a Fisher's exact test and t-test for categorical and continuous variables, respectively

Abbreviation: SD, standard deviation

**Table S2. Smoking EWAS meta-analysis results (unadjusted P-value < 0.05) in NAc, combined with ancestry-specific EWAS results.**

**Uploaded as a separate file due to size*

**Table S3. Smoking EWAS meta-analysis results in NAc.**

**Uploaded as a separate file due to size*

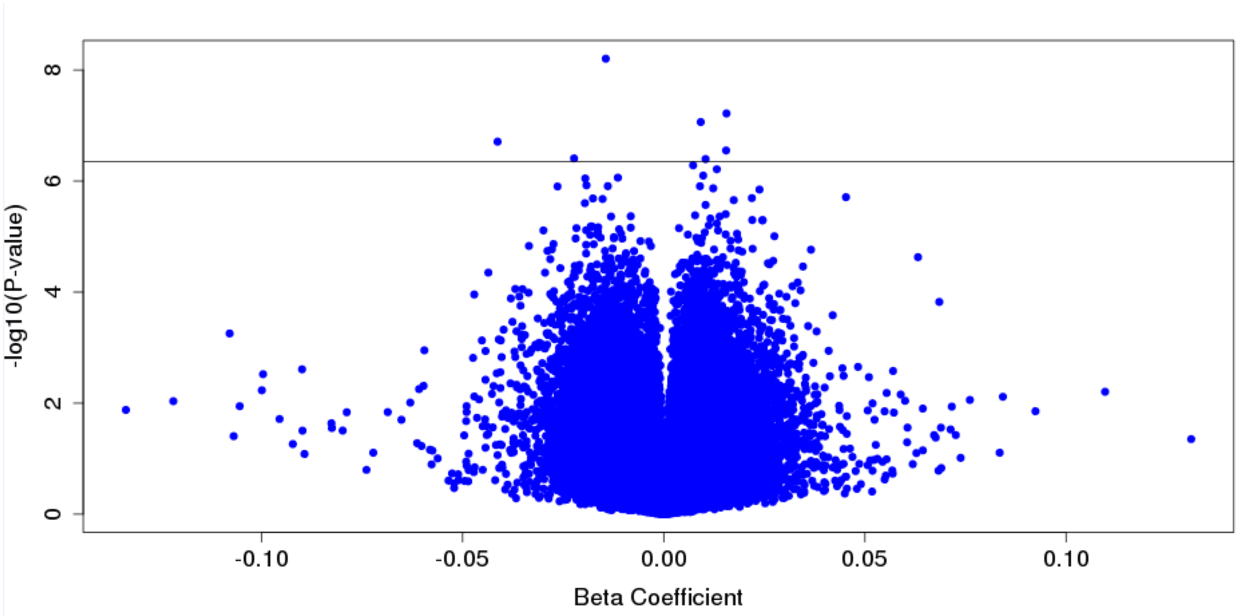


**Figure S1.** Smoking EWAS in NAc volcano plot. The solid horizontal line indicates genome-wide significance based on FDR<0.05.

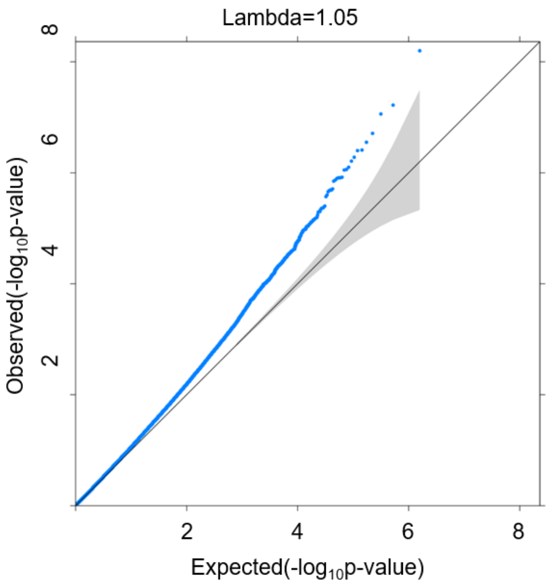


**Figure S2.** Quantile-quantile (Q-Q) plot of the smoking EWAS in NAc.

**Table S4. Top DNAm–RNAexp association for each of the NAc-identified smoking-associated CpG probes[1]**

| CpG | Chr | Position[2] | N genes tested in 1Mb window | Top DNAm–RNAexp association | | |
|---|---|---|---|---|---|---|
| | | | | β | P-value[3] | RNAexp gene tested |
| cg18384794 | 5 | 148509515 | 36 | 9.62 | $1.66 \times 10^{-3}$ | *ENSG00000283653* |
| cg17884843 | 3 | 147141132 | 10 | -5.65 | 0.08 | *ENSG00000231213* |
| cg05224975 | 20 | 57049253 | 25 | 2.70 | $9.52 \times 10^{-3}$ | *ENSG00000124222* |
| cg09959332 | 10 | 81224116 | 38 | 4.47 | $2.08 \times 10^{-2}$ | *ENSG00000214695* |
| cg11912754 | 8 | 48676898 | 14 | 2.28 | $9.69 \times 10^{-3}$ | *ENSG00000269924* |
| cg13555116 | 13 | 25862332 | 26 | 3.78 | $4.39 \times 10^{-2}$ | *ENSG00000273167* |
| cg08395748 | 16 | 67605701 | 97 | **4.03** | $\mathbf{1.54 \times 10^{-4}}$ | ***ENSG00000270049*** |

[1]All genes within 1 Mb of the smoking-associated CpGs and expressed (genes that did not have more than 10 counts in greater than 10% of the samples were removed) in NAc were tested

[2]CpG position based on GRCh37/hg19 human assembly

[3]P-value shown in bold met a Bonferroni correction for the number of genes tested by CpG (e.g., a significance threshold of $5.15 \times 10^{-4}$ [0.05/97] was applied for cg08395748)

Abbreviations: Chr, chromosome

**Table S5. Overlap between smoking-related CpGs in blood (FDR<0.05) and NAc samples.**

**Uploaded as a separate file due to size*

**Table S6. Overlapping genes with smoking-associated DNAm changes identified in blood. Only blood-based smoking-associated CpGs (FDR<0.05) annotated to an overlapping gene are presented.**

| Overlapping Gene[1] | CpG[2] | Distance to gene[3] | Blood-based EWAS meta-analysis[4] | | | |
|---|---|---|---|---|---|---|
| | | | Effect | SE | P-value | Dose-Response[5] |
| *ZIC1* | cg08013557 | 0 | 0.003 | 0.0006 | **$7.41\times10^{-6}$** | No |
| *ZIC1* | cg03631131 | 0 | 0.003 | 0.0009 | **$4.65\times10^{-4}$** | No |
| *ZIC1* | cg16209664 | 0 | 0.003 | 0.0008 | **$1.80\times10^{-3}$** | No |
| *ZIC1* | cg21657087 | 0 | 0.003 | 0.0009 | **$3.20\times10^{-4}$** | No |
| *ZCCHC24* | cg20618826 | 0 | -0.003 | 0.0010 | **$1.19\times10^{-3}$** | No |
| *ZCCHC24* | cg04920385 | 0 | -0.004 | 0.0010 | **$5.72\times10^{-5}$** | Yes |
| *ZCCHC24* | cg24501831 | 0 | -0.004 | 0.0010 | **$6.49\times10^{-6}$** | Yes |
| *ZCCHC24* | cg24000860 | 1,054 | -0.002 | 0.0007 | **$5.37\times10^{-4}$** | Yes |
| *ZCCHC24* | cg22653140 | 8,843 | -0.003 | 0.0011 | **$1.90\times10^{-3}$** | No |
| *PRKDC* | cg26982601 | 8,261 | -0.005 | 0.0014 | **$6.26\times10^{-4}$** | No |
| *PRKDC* | cg02895394 | 0 | -0.006 | 0.0016 | **$6.98\times10^{-5}$** | Yes |
| *PRKDC* | cg01286191 | 0 | -0.003 | 0.0007 | **$2.08\times10^{-5}$** | Yes |
| *PRKDC* | cg27320734 | 0 | -0.004 | 0.0009 | **$1.82\times10^{-5}$** | Yes |
| *PRKDC* | cg17496794 | 0 | -0.004 | 0.0010 | **$7.03\times10^{-6}$** | Yes |

[1]Blood-based smoking-related CpGs (N=18,760, FDR<0.05) were annotated to the nearest GENCODE v29 gene within a 1 Mb window. In total, 21,730 genes were annotated to a nearby CpG, of which 10,408 were unique. Similarly, we annotated NAc-based smoking-related CpGs (N=7, FDR<0.05) to the nearest gene within a 1 Mb window. These 3 genes showed evidence of differential methylation by smoking in blood and NAc

[2]Smoking-associated CpG in blood-based EWAS meta-analysis (FDR<0.05; Joehanes R, et al. 2016) annotated to overlapping gene

[3]Minimum distance (bp) from the CpG to the gene body

[4]Published results pulled from Supplemental Table 2 or 3 from Joehanes R, et al. 2016

[5]Exhibit dose-response relationship (pack-years) at FDR<0.05 (published Supplemental Table 3)

Abbreviations: SE, standard error

**Table S7. Blood-based SmPEGS1 applied in NAc samples.**

| Samples | OR (95% CI)[2] | P-value[2] | AUC (95% CI) |
|---|---|---|---|
| EA + AA | 1.78 (1.22, 2.59) | **0.003** | 0.62 (0.54, 0.71) |
| EA | 2.00 (1.12, 3.58) | **0.020** | 0.67 (0.55, 0.78) |
| AA | 1.45 (0.79, 2.67) | 0.236 | 0.55 (0.43, 0.67) |

[1]Sugden, *et al.* 2019
[2]Based on a logistic regression model, adjusting for same set of covariates as in the EWAS
Abbreviations: SmPEGS, smoking methylation PolyEpigenetic Score; EA, European American; AA, African American; SE, standard error; OR, odds ratio; CI, confidence interval; AUC, area under the receiver operating characteristic curve
